# The impact of taxonomic change on the Amazonian palm flora

**DOI:** 10.1101/2024.11.29.625979

**Authors:** Juliana Stropp, Andreza S. S. Pereira, Thaise Emilio, Leila Meyer, Rafaela Trad, Fernanda Alves-Martins, Richard J. Ladle, Joaquín Hortal

**Author notes:** **Author for correspondence:** Juliana Stropp.

## Abstract

Although taxonomy is an evolving discipline, taxonomic change is rarely accounted for in macroecological studies. By tracking the history of species descriptions and synonymizations of more than 800 names of Amazonian palms, we reveal shifts in species counts across space and time, the factors associated with taxonomic lumping, and the time needed to detect synonyms. The 175 Amazonian palm species known to date are the result of a gradual accumulation of new descriptions and a few distinct revisions events that led to the recognition of approximately 800 heterotypic synonyms. Most of these synonyms were detected after the mid-1990s and caused an 80% decrease in species counts within just three decades. The time to detect synonyms ranged from 3 to 227 years. Species with a large population, widespread distribution, and those described early were more frequently lumped. This suggests undetected synonyms inflate species richness unevenly across taxa and regions. Our study highlights how much taxonomic revisions can affect our understanding of biodiversity. Without them, we risk building our ecological analyses and conservation assessments on shaky ground.

## Introduction

About four thousand new plants names are proposed each year. However, only half of these result from new field discoveries; the other half stems from the taxonomic revision of already described species [1]. Such revisions often result in lumping, splitting, or reassigning species to a different genus. When substantial, these changes can alter species counts across taxa and reshape the geographic patterns of species richness [2]. Yet, their impact on biodiversity patterns remains largely unquantified. This knowledge gap is particularly problematic for hyper-diverse ecosystems like the Amazon rainforest, where new species discoveries and revisions are common [3] and may even affect estimates of global species richness [4].

Taxonomic lumping and reassignments are frequent in many plant groups [5]. Taxonomic lumping happens when species boundaries are re-evaluated and modified, resulting in combining two or more previously accepted species into a single one. Reassignments, on the other hand, occur when relationships among species are reinterpreted, resulting in species being moved from one genus to another. Both, lumping and reassignments, generate synonyms, i.e.,, duplicated names for the same taxon (see box 1 and table S1 for definitions). Overall, synonyms tend to outnumber the counts of accepted vascular plant species by a factor of 1.5 [6]. The number of synonyms across flowering plants (angiosperms) is not homogeneous but it varies by several orders of magnitude across taxa. Half of the angiosperm species are not associated with a single synonym (i.e., have only a single accepted name assigned to their original description), whereas a few angiosperms have been accumulating up to 300 synonyms [7]. The identification of synonyms may take only a few years or up to centuries after the first proposal of a name [8].

Yet, our understanding of the impacts of taxonomic change on plant species richness is mainly derived from theoretical models [9] or from other biological groups, such as birds or amphibians [10]. Complementing the theoretical work with empirical studies has long been hampered by limited data availability [11]. However, a growing volume of digital resources, including botanical taxonomic treatments (e.g., monographs, revisions, and synopses), and easy access to plant nomenclatural databases [e.g., 12,13] opens new opportunities to explore changes in species taxonomy. Such data, to our knowledge, have not been used yet to assess how taxonomic change has shaped the current pattern of plant richness at a broad temporal and spatial scales.

Here we analyse 200 years of taxonomic descriptions and lumping for all Amazonian palms to uncover the changes in the number of accepted species across genera and regions. Our study addresses three questions: 1) How did the counts and proportion of accepted species vary over time across Amazonia? 2) How long can it take to recognize an accepted species as a synonym? and 3) Which factors are associated with taxonomic lumping?

We chose Amazonian palms as a model group because they are among the best-studied groups of the tropical flora [14], with a wealth of species descriptions and revisions published since the 18^th^ century [14]. Together with contemporary field sampling [15], this provides rich baseline data for pertinent analyses. We also think that a deeper understanding of taxonomic progress in Amazonia is needed. In times of climate change, global mass extinction, and decreasing research budgets for taxonomy, we risk losing many species before they are adequately described or classified [16]. Thus, we discuss the implication of changes in species taxonomy for macroecology and biodiversity conservation analyses that rely on species count and identity.

## Materials and methods

### Establishing a checklist of Amazonian palms and associated synonyms

We considered as Amazonian palms all palm species that occur at elevations below 500 m a.s.l. and within the biogeographic limits of Amazonia [17]. To create a broadly accepted checklist of Amazonian palms, we included only species (excluding hybrids, varieties, and subspecies) listed in any of the four checklists: Henderson (1995) [18], in Cardoso et al. (2017) [19], ter Steege *et al*. (2019) [3], and Flora e Funga do Brasil (2020) [20]. Differences between these four lists reflect variations in the geographical boundaries of Amazonia and taxonomic interpretations of what constitutes an accepted species or a synonym. Our initial broadly accepted checklist contained 240 species (table S1).

We then verified the taxonomic status of the 246 species names against four taxonomic data portals: Tropicos [21], Plants of the World Online (POWO) [22], World Flora Online (WFO) [23], and Flora e Funga do Brasil [20]. For all names identified as synonyms or unplaced in any of the data portals, we obtained the respective accepted name. This standardization resulted in a checklist of 206 accepted species names. All names were found at least in one data portal; the unique identifiers provided by individual data portals are given in table S1.

Out of these 206 species, 53 were not included in Henderson [18], which is the most recent checklist of Amazonian palms compiled by an Arecaceae specialist. We thus verified the geographical range of these 53 species in Flora do Brasil [20]; “Palms of the Americas” [24], the website ‘Palmweb’ [25], and the pertinent taxonomic literature. This cross-referencing identified 31 species that do not seem to occur below 500 m a.s.l within the biogeographic limit of Amazonia. These species were excluded from our analyses. Therefore, our final checklist includes 175 accepted palm species, described until 2012 (the last year with consolidated data in the checklists of Cardoso et al. [19] and ter Steege et al. [3]). Our list comprises species of different sizes and growth forms, e.g., large tall-stemmed palms, large to small acaulescent palms, small palms, and climbing palms [26].

We also retrieved the following information from the four taxonomic portals: (1) a list of homotypic and heterotypic synonyms (see Box 1) associated with the 175 currently accepted palm species; and (2) ancillary bibliographical information (i.e., author, title of the taxonomic work, and year of publication) for the currently accepted species and their synonyms. We manually checked all names listed as synonyms (N = 1285) in the four taxonomic portals. We corrected obvious misspellings and flagged names that could not be assigned to a known species, as they were illegitimate or not validly published designations (table S1). We also flagged names that we could not associate with a publication or which were not available in the database of the International Plant Names Index [27]. Such manual checks were necessary because information on synonyms retrieved from the four taxonomic portals may be inconsistent.

We then selected only heterotypic synonyms, which result from merging previously accepted species that were described based different type specimens (Box 1). Heterotypic synonyms were identified by querying the Plants of the World database [12] with the R package ‘kewr’ [13] and by consulting taxonomic works, in which the currently accepted species have their synonyms (or unplaced names) mentioned. Finally, we manually checked the earliest taxonomic work, including monographs, synopses, and revisions, to determine the year of synonymization for all heterotypic synonyms. These taxonomic works were identified through searches of main virtual botanical databases, such as Tropicos [21], POWO [22], and WFO [23]. As our focus is on temporal trends in the number of accepted palm species, we considered only heterotypic synonyms that are recognized at species level. This is justified because species is the analytical unit for most (macro)ecological and conservation studies, we thus excluded subspecies, hybrids, and varieties from our analyses.

#### Box 1.

**Definition and examples of homotypic and heterotypic synonyms**

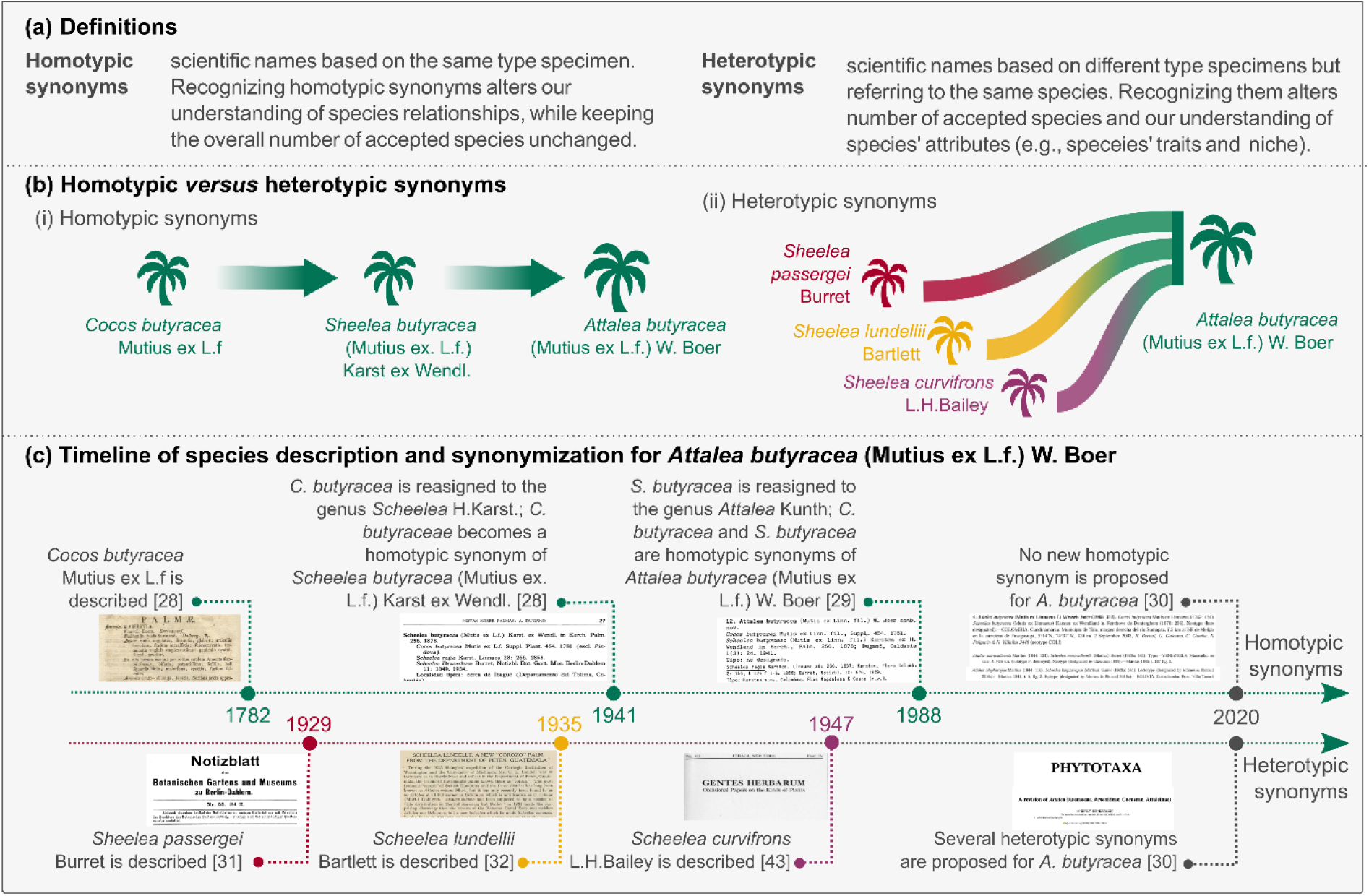

### Herbarium records of Amazonian palms

We assessed the spatial variation in the proportion of accepted species in all described taxa (accepted species plus heterotypic synonyms) varies across Amazonia. For this, we used the location of 15,446 herbarium records of all 175 species of Amazonian palms, which were retrieved from the Global Biodiversity Information Facility [36]. We screened all records and flagged those with uncertain geographical coordinates and/or missing taxonomic information at species level. This filtering led to a dataset of 8249 herbarium records for 170 palm species. The ‘rgbif’ R package [37] was used to retrieve records, and the ‘bdc’ R package [38] to flag records with potential errors.

### Data analysis

#### Temporal trends in the proportion of accepted species

To analyse temporal trends, we followed Alroy [8] and calculated for each year *n* the proportion of accepted species (*P*) in all described taxa as:

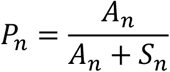

where *A* stands for the number of accepted species and *S* denotes the number of heterotypic synonyms in year *n*. To visualize the time needed to recognize a previously accepted species as a heterotypic synonym, we used Kaplan-Meier curves [39]. Such curves are commonly used in medical research to visualize the probability of an event (e.g., death) over time for different treatment groups. Here, we considered the year of taxonomic description as the base year of the observation period (*t*_*0*_) and the year of synonymization as the year in which a taxonomic event is recorded (*t*_1_), and from which onwards a name no longer corresponds to an accepted species. The Kaplan-Meier curves were established considering only heterotypic synonyms. Both the proportion of accepted species in all described taxa and the Kaplan-Meier curves were calculated for: (1) all taxa; (2) taxa grouped cohorts according to year of description (i.e., until 1900; between 1901 and 1950; between 1951 and 1980; and between 1981 until 2012); and (3) taxa belonging to the three most species-rich palm tribes (Cocoseae, Euterpeae, Geonomateae). The Kaplan-Meier curves were established with the ‘survival’ [40] and ‘survminer’ [41] R packages.

#### Spatial trends in the proportion of accepted species

To assess whether species have a higher number of heterotypic synonyms in certain regions of Amazonia, we mapped the proportion of accepted species in all described taxa. A lower proportion of accepted species in a given area may indicate a higher number of heterotypic synonyms, suggesting a higher number of duplicated descriptions for the same species. This approach allows us to identify spatial patterns in synonymy across the region. For this, we used a grid of resolution of 100 × 100 km, which covered the Amazonian region [17] and was projected to the ‘South America Albers Equal Area Conic’ (crs = “ESRI 102033”). Based on herbarium records retrieved from GBIF, we calculated for each grid cell the proportion of accepted species relative to the total number of names (i.e., accepted species plus heterotypic synonyms). We also determined the average proportion of accepted species for all cells, within latitudinal and longitudinal bins of 100 km. The map of proportion of accepted species was obtained for two distinct periods of time: (1) from 1769 to 1990, and (2) from 1769 to 2020. We chose these periods because they align with substantial changes in the total number of accepted palm species in Amazonia.

#### Correlates of taxonomic lumping

We tested whether taxonomic lumping has been more frequent for species that occupy large areas, were described in early years, and have a large population. To do so, we tested for correlations between: (1) number of heterotypic synonyms; (2) year of first taxonomic description; (3) size of the species geographic range; (4) number of individuals; and (5) estimated population size for each accepted species. Pearson correlation coefficients were calculated using the function *corphylo* in the *‘ape’* R package [42]. The phylogenetic relationship among species was extracted from the Maximum Clade Credibility phylogenetic tree for palms proposed by Onstein *et al*. [43]. These correlations reflect to some extent the interplay between species characteristics and taxonomic revisions, as taxonomic lumping arguably expands species’ geographic ranges and their population size.

The geographic range was calculated based on range maps for Amazonian palms, which were obtained with the function *BIEN_ranges_load_species* from the *‘BIEN’* R package [44]. Range maps were re-projected to South America based on Albers Equal Area Conic projection. We then calculated the area (km^2^) of polygons representing the known range of a species. We considered the entire geographic range of each species, encompassing Amazonia and other biomes in the Neotropics. As these maps are based on collection localities of herbarium vouchers, their accuracy depends on the number of vouchers available [44]. Range maps was available for 155 species of the 175 species in our dataset.

The observed and estimated population size were obtained from ter Steege et al. [15]. Observed population size represents the abundance of individual species observed in 1170 tree inventory plots established across Amazonia, whereas the estimated population size was obtained by modelling species abundance and density for Amazonia [see 15 for details]. Observed and estimated population size was available for 57 species of the 175 species in our dataset.

## Results

Our dataset contains 1460 unique names, including 175 accepted species and 1285 names that comprise both homotypic and heterotypic synonyms, as well as unplaced names or designations (e.g., orthographic variants, illegitimate names, or *nomina nuda*). Of the 175 accepted species, 68 had no synonyms or only homotypic synonyms, while 107 were associated with a total of 952 heterotypic synonyms. For 940 of these heterotypic synonyms, we were able to retrieve both the year of publication and the year of synonymization. Among the heterotypic synonyms, 136 represented varieties or subspecies, and 804 were species-level heterotypic synonyms linked to 102 accepted species (Appendix S2). We successfully obtained the year of publication and the year of synonymization for 798 of the 804 species-level heterotypic synonyms (table S2). For brevity, we refer to these 804 species-level heterotypic synonyms as heterotypic synonyms.

### How did the counts of accepted species vary over time across Amazonia?

Our results show that species counts can vary significantly before and after a major taxonomic revision. The current number of 175 accepted species for the Amazonian palm flora is comparable to the count in the 1850s (Fig 1a), with the difference that in the 1850s there was not a single heterotypic synonym registered, whereas now there are 804. This indicates that the gradual accumulation of new species descriptions over two centuries has been compensated by taxonomic lumping, most of which occurred after 1990.

**Figure 1.**
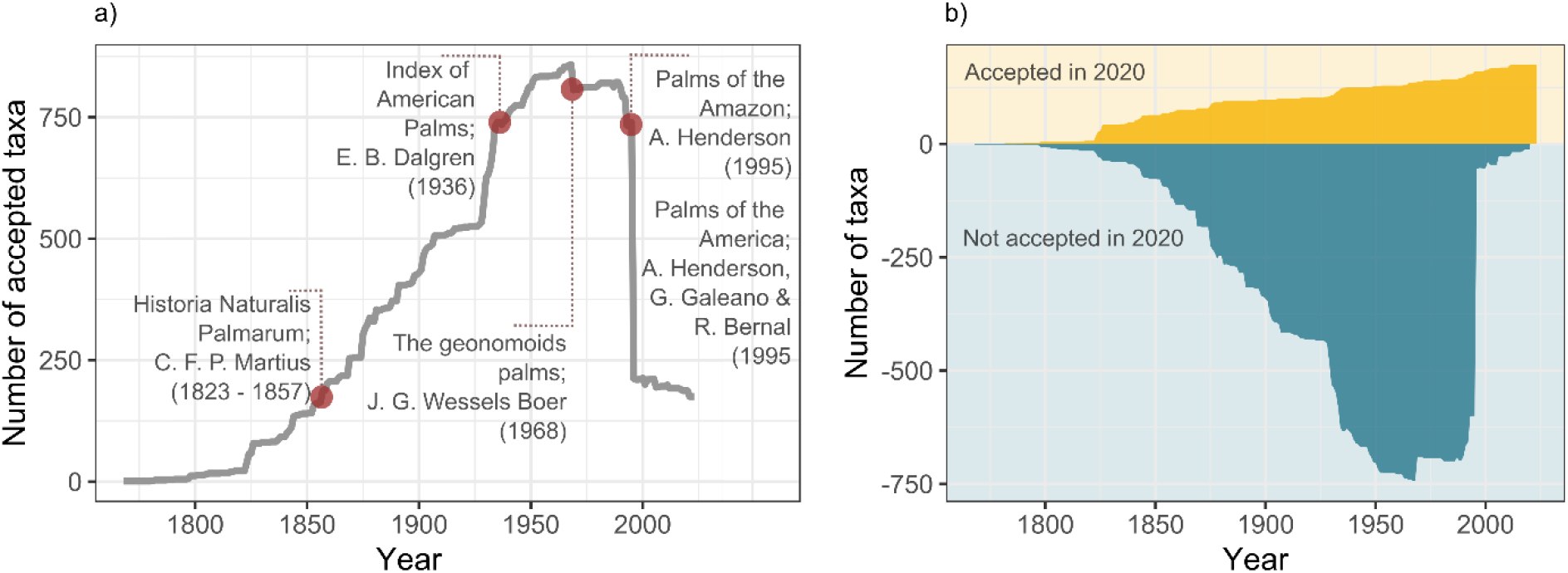
Timeline of accepted species for the Amazonian palm flora. a) Number of accepted taxa over time, red dots mark the year of major taxonomic works; b) Number of described taxa per year that are accepted (yellow) *versus* not accepted (blue) in 2020.

The number of Amazonian palm species increased steadily from the late 1700s until the mid-1960s and reached its maximum in 1966 when 854 taxa were recognized as distinct species. After nearly two centuries of adding almost continuously new descriptions, taxonomic lumping reversed this trend and led to an 82% decline in the overall number of accepted species in just three decades (figure 1a). The first significant drop, from 854 to 802 species, occurred in 1968 due to the work of Jan Gerard Wessels Boer [31]. The second drop was more substantial, with the number of accepted palm species declining by 79% from 787 in 1990 to 206 in 1995. This reduction stemmed mainly from synonymizations proposed by Henderson [18] and Henderson et al. [24]. The third and most recent drop in the number of accepted species (from 206 to 175) occurred in 2020, and coincided with the lumping proposed for *Attalea* Kunth [32]. In brief, counting Amazonian palm species in the mid-1990s and in 2020 would yield two very different totals, 731 and 175 respectively, which differ by a factor of 4.5 despite being only three decades apart (figure 1b). This striking observation underscores how critical taxonomic revisions are for obtaining accurate species counts. It also shows that taxonomic revisions can have significant knock-on effects on macroecological analyses, conservation assessments, and any analyses that rely on species identity and counts.

Of the 34 Amazonian palm genera, 11 reached their current species number in the 1990s and another 14 in the 2000s (figure 2a). *Attalea, Iriartella* H.Wendl. and *Syagrus* Mart. were the latest to reach their current number of accepted species, doing so in 2020. In contrast, *Raphia* P.Beauv. and *Wettinia* Poepp. ex Endl. were the earliest, reaching their current total before the 1900s. As the abundance of species can vary across geographic regions, different regions may reach their current number of accepted species at different times. We find that current species numbers were reached recently across most of Amazonia, with the exception of the central Amazon (figure 2b). In this region, frequently recorded genera had already reached their current number of accepted species around 1940.

**Figure 2.**
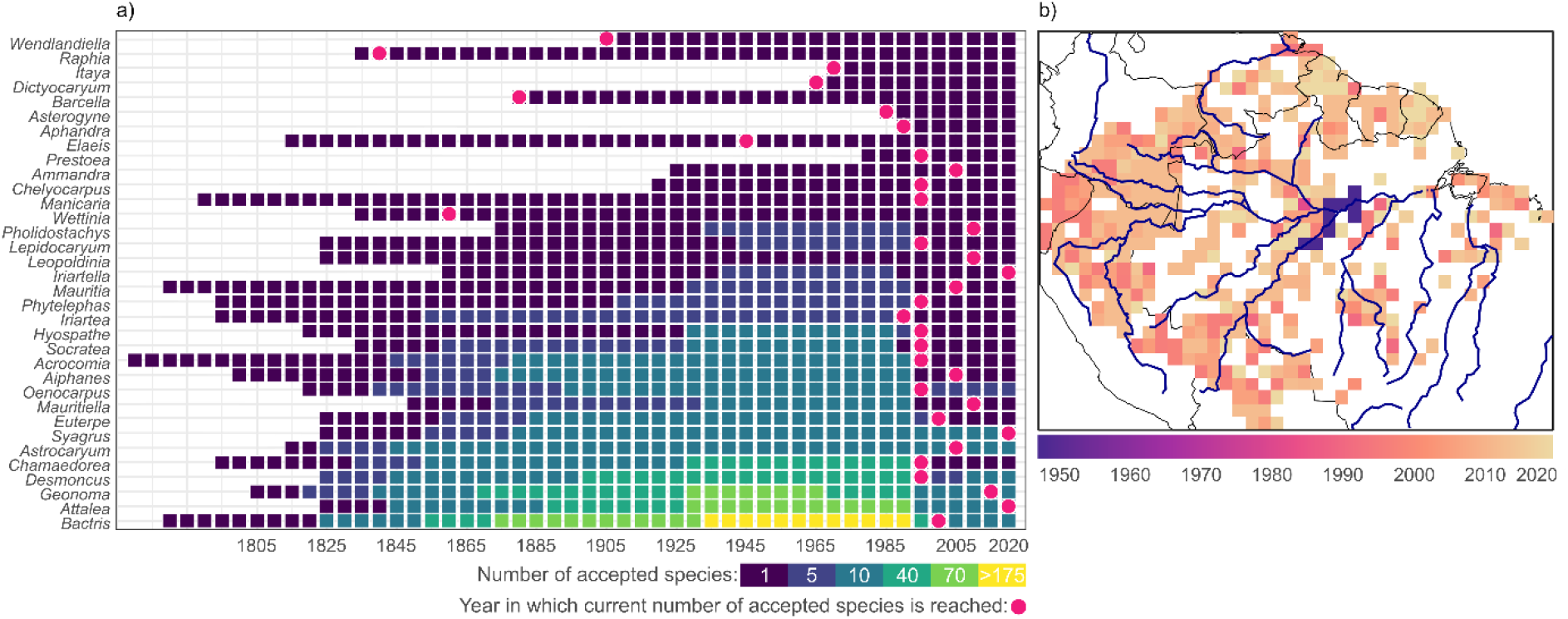
Timeline of accepted species across genera in Amazonia. a) Number of accepted species by genus over time; pink dots mark when the current number of accepted species was reached, blue to yellow squares indicate the number of accepted species. b) Map depicting the median year in which the genera recorded in each grid cell reached their current number of accepted species, with light colours indicating more recent years.

### How did the proportion of accepted species vary over time across Amazonia?

The proportion of accepted species in all described taxa mirrors the trend in taxonomic lumping described above. Overall, this proportion has remained relatively stable over 200 years and only started to decrease in the late 1960s. However, a more detailed examination suggests that lumping is less frequent in recently described species than those described earlier (Figs. 3a and 3b). Nearly 80% of the species described after the 1990s are still considered accepted to date. This share is only 40% for taxa described between 1950 and 1980. Among the three most species-rich tribes, Geonomateae has the highest share of species that remain accepted (figure 3c).

**Figure 3.**
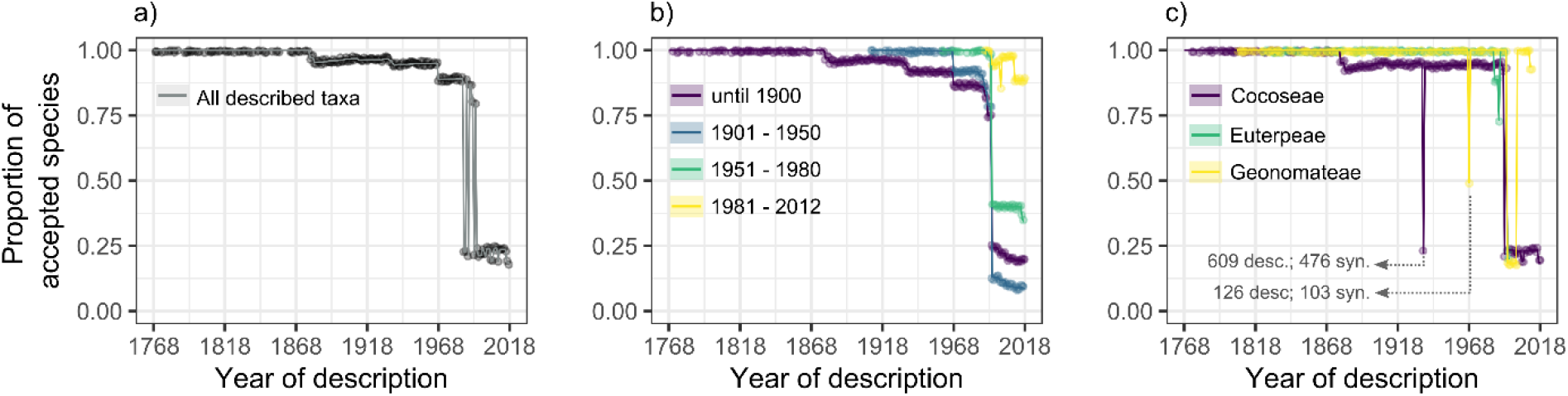
Temporal decay in the proportion of accepted species. (a) all taxa (oscillating grey line results from large changes in the proportion of accepted species after the 1990s); (b) all taxa grouped into cohorts based on the year they were described; (c) taxa belonging to one of the three most species-rich palm tribes (Cocoseae, Euterpeae, or Geonomateae)

Considering the spatial context, in 1990, the proportion of accepted species in described taxa was rather homogeneous and close to 1 across the entire Amazon (figure 4a). However, in the 2020s, this proportion decreased to between 0.1 and 0.4 throughout the entire region (figure 4a and 4b). A comparison of two maps for 1990 and 2020 reveals that species that are typically found in Western Amazonia are associated with a particularly high number of heterotypic synonyms. Relevant genera include *Attalea, Bactris* and *Geonoma* Willd., which reached a particularly high number of synonyms after the 1990s (Fig 4c and 4d). These results show that taxonomic revisions are not homogeneous across space, time, and taxonomic groups. Rather they occur at distinct moments in time with substantial impacts that vary across regions and genera.

**Figure 4.**
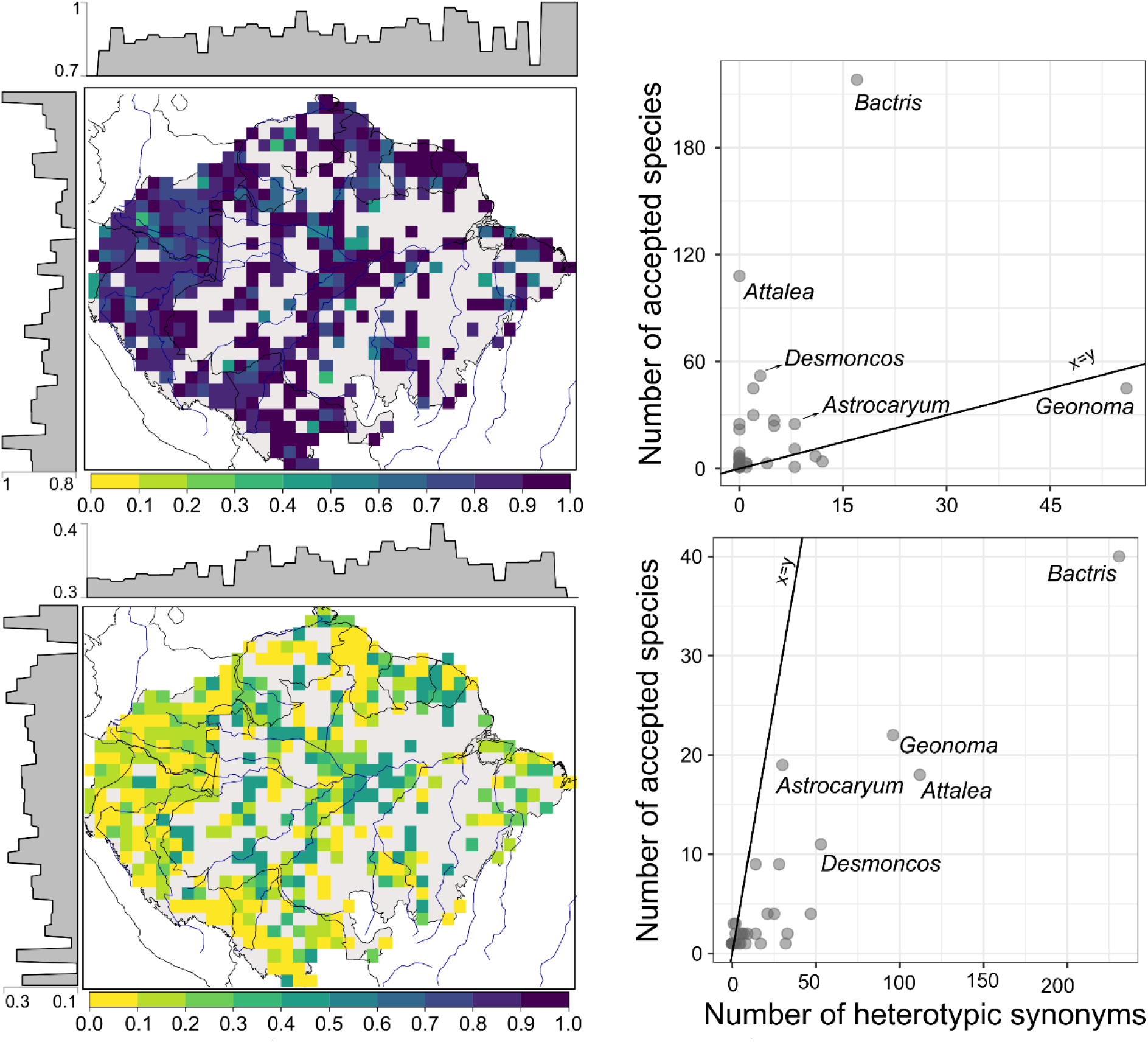
Proportion of accepted species in all described taxa. (accepted species and heterotypic synonyms). Maps depict proportions for 1990 (a) and 2020 (b). Lighter shades represent a lower proportion of accepted species (i.e., more heterotypic synonyms per accepted species), while darker shades represent a higher proportion. Histograms show the mean proportion of accepted species for each longitudinal and latitudinal band of 0.9 degrees (100 km). Scatterplots show the number of accepted species versus heterotypic synonyms recognized in 1990 (c) and in 2020 (d); black line indicates a 1:1 ratio.

### How long can it take to recognize an accepted species as a synonym?

For Amazonian palms, it took anything from 3 to 227 years to detect a heterotypic synonym (median = 79 years). The time required to detect synonyms depends on the age of the taxonomic descriptions and the tribe to which the species belongs to. It has taken 79 years to detect about half of all currently recognized heterotypic synonyms. An additional 147 years was required to detected the other half (table 1; figure 5a). For taxa described before the 1900s, it has taken on average 114 years to identify half of them as synonyms and a total 220 years to detect all synonyms (figure 5b). The detection time is shorter for more recently described taxa. For example, it only took an average of 26 years to detect synonyms among palms described between 1951 and 1980, and 18 years for those described between 1991 and 2012. The time lag between the first taxonomic description and synonymization also varies depending on the palm tribe. Among the top three tribes in terms of number of species, Euterpeae and Geonomateae had half of their synonyms detected approximately 60 years after description. In contrast, Cocoseae had half of the synonyms detected on average 90 years after description (figure 5c). The average time to detect heterotypic synonyms within each tribe ranged from 60 (Lepidocaryeae) to 164 years (Leopoldinieae). This result shows that there are prolonged periods without taxonomic revisions. During these periods, species counts can be highly uncertain, as potential heterotypic synonyms remain “hidden” among the accepted species.

**Table 1.** Summary statistics of Kaplan-Meir curves, indicating the time to detect heterotypic synonyms of Amazonian palms.

|  | **N heterotypic synonyms** | **Time to detect 50% of all heterotypic synonyms** | **Time to detect the first 10% of all heterotypic synonyms** | **Time to detect the last 10% of all heterotypic synonyms** |
| --- | --- | --- | --- | --- |
| All heterotypic synonyms | 798 | 79 | 38 | 138 |
| Taxa described until 1900 | 422 | 114 | 84 | 152 |
| Taxa described between 1901 and 1950 | 327 | 61 | 40 | 67 |
| Taxa described between 1951 and 1980 | 34 | 26 | 7 | 32 |
| Taxa described between 1981 and 2012 | 15 | 18 | 10 | 18 |
| Cocoseae | 505 | 90 | 5 | 138 |
| Euterpae | 56 | 63 | 44 | 125 |
| Geonomateae | 101 | 65 | 29 | 138 |

**Figure 5.**
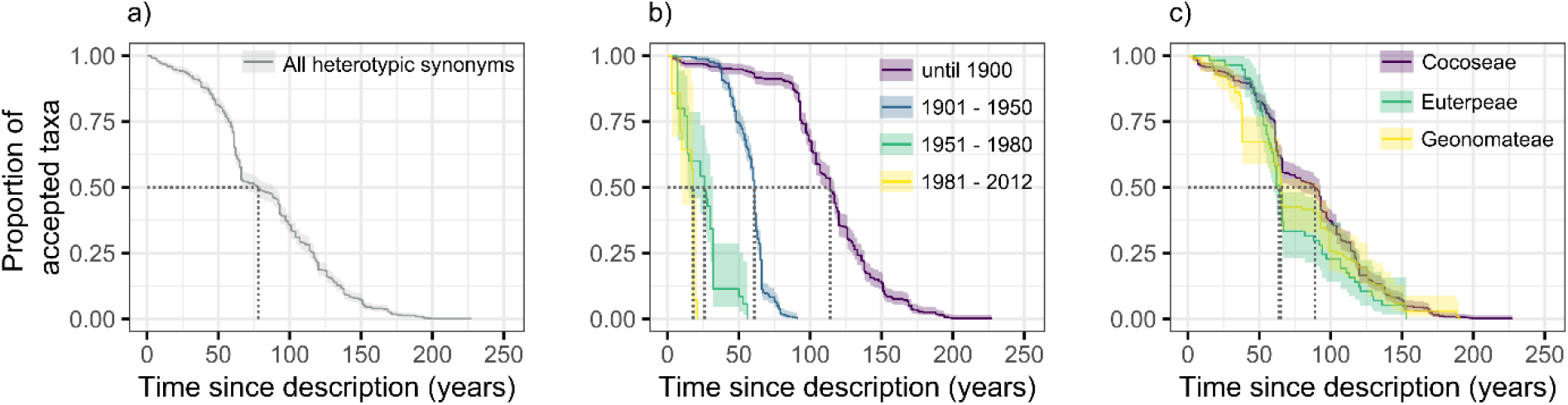
Time-lag between species description and synonymization. Kaplan-Meier curves for (a) all heterotypic synonyms, (b) by cohorts according to year of taxonomic description, and (c) heterotypic synonyms associated with accepted species belonging to Cocoseae, Euterpeae, or Geonomateae. Curves reach zero (y-axis) because only heterotypic synonyms were considered in this analysis.

### Which factors are associated with taxonomic lumping?

Species that were early-described, are widespread and have a large population tend to have a higher number of heterotypic synonyms (Table2). This finding may indicate that such species are more subject to taxonomic lumping. However, it could also point to a reversed causality through which species reach a large geographic range and population size simply as a result of recurrent taxonomic lumping. Additionally, we found that widespread and abundant species tend to be described earlier (Table 2).

**Table 2.** Pearson correlation coefficients and significance levels (\*\*\**p* < 2.2e-16; \*\**p* < 0.001; \**p* < 0.01).

| Parameter | Number of heterotypic synonyms | Size of area of occupancy | Year of description | Number of individuals | Population size |
| --- | --- | --- | --- | --- | --- |
| Number of heterotypic synonyms | 1 |  |  |  |  |
| Size of area of occupancy | 0.6*** | 1 |  |  |  |
| Year of description | -0.2*** | -0.5*** | 1 |  |  |
| Number of individuals | 0.4** | 0.4*** | -0.4** | 1 |  |
| Population size | 0.2** | 0.4*** | 0.01* | 0.9*** | 1 |

## Discussion

Our study shows that the impact of taxonomic progress on species richness is substantial but not homogeneous across time, space, and palm taxa. Over the past two and a half centuries, taxonomic descriptions of Amazonian palms have steadily accumulated, while lumping occurred through distinct and comprehensive taxonomic revisions published mostly after 1990 [18,24]. As a result, more than 800 heterotypic synonyms remained unrecognized for periods ranging from a few years to more than two centuries. Lumping has been recurrent for species described in early years, that occupy a large area and have a large population. Moreover, species that typically occur in Western Amazonia were subject to more taxonomic lumping than species that occur elsewhere in Amazonia. Currently, accepted species represent only 12% of all described names. The observed taxonomic lumping might even be, to some extent, excessive because some palm lineages might harbour cryptic species [45]. Our results show that taxonomy is far from static. Palms have benefited from a well-connected community of taxonomists and ecologists, and a species-level phylogeny will soon be completed [46]. Thus, future progress will continue to refine species boundaries as new data and methods emerge. We think, however, that further extensive lumping of Amazonian palms appears unlikely. Next, we discuss how shifts in species number and identity due to taxonomic changes may influence macroecological and conservation analyses.

### Taxonomic progress and its impact on the Amazonian palm flora

The current Amazonian palm flora, comprising 175 accepted species, resulted from a gradual accumulation of species descriptions and distinct events of taxonomic lumping through which *ca*. 800 heterotypic synonyms were detected. Thus, the Amazonian palm flora has not been growing continuously through the accumulation of new species descriptions. It is also not the result of a ‘zero-sum game’ through which increases in species numbers from descriptions of new species are directly balanced by reductions in species numbers caused by taxonomic lumping (Fig 1b). Instead, new species were added, and previously accepted species were refuted unevenly across taxa, geographic regions, and time.

Changes in species taxonomy arise from evolving taxonomic practices [47]. Before the 1950s, species-level taxonomy for palms was in a “state of chaos” as remarked by Henderson and colleagues (1995) [24]. Early taxonomists, such as J. Barbosa Rodrigues (1842 - 1909), L. H. Bailey (1858 – 1954), and C. E. M. Burret (1883 - 1964) accepted minimal morphological variation within species and described nearly every specimen as a new species. It was only in 1968 that J. Wessels Boer (1936 - 2019) initiated major synonymizations for Geonomoid palms (figure 1). In the 1970s, taxonomy shifted in Latin America from local, country-specific studies to a regional, monographic approach driven by the Flora Neotropica project. This initiative, involving taxonomists working across 30 countries [48], encouraged the study of morphological variation across broad geographic ranges and influenced subsequent taxonomic revisions, including those of Henderson and colleagues [18,24,32,32,32,49– 52]. For example, revisions of *Geonoma* (68 species) and *Attalea* (30 species) are based on the analyses of morphological characters of 4,990 and 902 specimens, respectively. In result, 96% of palm species described by C. E. M. Burret were synonymized between the 1960s and 2000s.

The recent lumping is not restricted to Amazonian palms. *Guatteria* Ruiz & Pav. (Annonaceae), for example, lost the status of the most species-rich genus of woody Neotropical trees after significant taxonomic lumping in 2015 [53]. Other groups of tropical plants experienced similar lumping. The large genus *Myrcia* DC. ex Guill. (Myrtaceae), includes several species that have been described multiple times and were later synonymized [54]. To date, *Myrcia guianensis* (Aubl.) DC., for example, is associated with approximately 150 heterotypic synonyms [55]. In this context, it is interesting that at global scale, the overall number of accepted plant species is still increasing [56]. This increase is well-documented and results mostly from new species descriptions [57]. However, the extent to which lumping offsets this growth is difficult to track and quantify because taxonomists are not required to explicitly record in the taxonomic literature (e.g., botanical monographs) the proposal of heterotypic synonyms [58].

For seed plants, it is estimated that accepted species account for 34% [59] to 40% [6] of all taxonomic descriptions for this group, indicating that more than half of the described taxa were at a later point synonymized. These numbers are roughly twice as high as the 21% observed here for Amazonian palms (considering only heterotypic synonyms). Multi-taxa comparisons of synonym rates, however, typically do not distinguish between homotypic and heterotypic synonyms [58,60]. While homotypic synonyms arise when a species is reassigned to another genus heterotypic synonyms result from changes in the delimitation of a given species. By explicitly distinguishing between these two types of synonyms when calculating synonym rates, we can improve our understanding of how changes in taxonomy, whether due to changes in delimitation or reassignments, affect species counts across taxa.

### Taxonomic lumping across taxa and regions

Taxonomic lumping has been more frequent for palm species that occupy large areas and have large populations. This finding is expected because taxa covering larger areas tend to show higher morphological variation due to a higher diversity of environmental conditions [61]. Particularly, flower morphology within species can vary substantially with climate, soil, and pollinator communities [62]. The resulting morphological plasticity can lead to independent taxonomic descriptions, which were likely prevalent in early years when the exchange of botanical information was limited and taxonomic studies had a restricted geographic scope. However, one could also argue that the causality is reversed. Both large phenotypic variation and broad range size could themselves be the consequence of taxonomic lumping. Therefore, our statistical analysis leaves room for interpretation and ambiguity regarding the actual causes and effects of taxonomic lumping.

Our data also show that western Amazonia harbours species that were subject to recurrent taxonomic lumping, reflecting a more complex taxonomic history in this region than elsewhere in Amazonia. This complexity may stem from the geo-political configuration of Western Amazonia, which includes several cross-country borders. Before the 1970s, the focus of botanical monographs on individual countries may have hindered exchange of botanical information and increased the likelihood of independent descriptions of the same taxa. However, many species that were later synonymized had been described based on specimens sent to C. E. M. Burret at the Herbarium Berolinense in Berlin (Germany) or to L. H. Bailey at the Hortorium Herbarium in Cornell (USA) [24]. Many of Burret’s type specimens were destroyed during World War II [51], complicating efforts to trace back the actual collection locations. As a result, it remains challenging to determine how taxonomic practices contributed to an over-description of species in western Amazonia.

An alternative explanation for the high number of heterotypic synonyms in Western Amazonia is that the region comprises recently diverged lineages that diversified within the region’s mosaic of habitat and fertile soils [63]. These lineages, initially described as distinct species, underscore the difficulty of defining species boundaries for closely related groups [47]. This suggests that the higher than expected species richness of palms in Western Amazonia [64] might need to be re-evaluated using post-2007 taxonomic revisions—such as updates for *Attalea* [32], *Bactris* [50], *Desmoncus* [49], and *Geonoma* [52]. Such research could determine whether the excess palm richness in Western Amazonia is caused by ecological and evolutionary processes or to some extent by taxonomic artefact.

If taxonomic lumping is not random across taxa and regions – as we observe here – and taxa show a strong spatial phylogenetic structure [65], then regions have reached their current number of accepted species at different points in times. Genera that typically occur in central Amazonia have reached their current number of species 50 years earlier than those occurring in other Amazonian regions. This observation may be influenced by the high frequency of *Elaeis oleifera* (Kunth) Cortés recorded in central Amazonia. *Elaeis* Jacq. is a genus with one single accepted species in the Americas [24] that is mostly recorded in central Amazonia [66]. By contrast, *Geonoma* and *Bactris*, which are recorded more frequently in Western Amazonia, are among the most species-rich genera of Amazonian palms and have a high number of heterotypic synonyms. It is important to note that the map depicting when genera reach their current number of species (figure 2b) is draw from digital herbarium specimens and thus is subject to incompleteness and biases. Palms are particularly underrepresented in herbaria, as their often large and spiny leaves are difficult to collect and transport [24]. Moreover, both palm inventories and taxonomic studies have been particularly abundant in western Amazonia in the last 30 years [67], which may have led to more species descriptions and detection of synonyms over time. Taken together, these observations highlight the importance of considering the status of taxonomic research when interpreting broad-scale patterns of species richness and turnover in species composition [68].

### Timing of taxonomic lumping

Our results show a large variation in the time heterotypic synonyms remain undetected, with recently described taxa being recognized as heterotypic synonyms more quickly than older ones. Moreover, taxa belonging to the tribes Euterpae or Geonomateae are recognized as heterotypic synonyms on average 100 years earlier than those belonging to the Cocoseae, which includes some of the most species-rich genera of Amazonian palms, such as *Attalea* and *Bactris*. The temporal variation arises because most synonyms were proposed in a few distinct taxonomic revisions: Amazonian palm flora in 1995 [18,24], *Bactris* in 2000 [50], *Geonoma* in 2011 [51], and *Attalea* in 2020 [32]. Thus, the time required to detect synonyms in Amazonian palms has been determined by how many years passed between the first species description and rather singular and comprehensive revision events.

These results suggest that heterotypic synonyms can remain hidden for long periods and may even remain unrecognised, given the limited taxonomic coverage in botanical monographs [69]. Yet, a detailed understanding of temporal trends in the detection of synonyms, especially for heterotypic synonyms, is needed for a broader range of taxa. A comparative analysis, particularly across tropical plants, could provide valuable insights into uncertainty on species counts and identity that may arise from the lack of revisionary work.

### Taxonomic lumping and hyperdominance in the Amazonian palm flora

Palms are among the most dominant species in the Amazonian tree flora [15]. The extent to which changes in species delimitation affect dominance patterns is subject to ongoing discussions [9]. If an accepted species resulted from the lumping of several species, this accepted species will have a larger population and a wider geographic range than before the lumping. Such a species could then also become hyperdominant. For palms we observed a moderate correlation (*r* = 0.4) between the number of heterotypic synonyms and the estimated population size, suggesting that taxonomic lumping, to some degree, may influence the current pattern of species dominance. Given the substantial lumping proposed for Amazonian palms, we expect that further lumping is unlikely to happen in the near future. Consequently, changes in species dominance may arise if future research uncovers cryptic species within large populations. Recent molecular evidence suggests that *Astrocaryum murumuru* Mart., with four heterotypic synonyms and an estimated population of 2.4 billion individuals [15], could potentially include such cryptic species [45]. This could justify a re-evaluation of species boundaries within this group. However, it is not possible to draw general conclusions about the relationship between hyperdominant species and taxonomic lumping or taxonomic splitting, which can reveal cryptic species. Both lumping and splitting depend on taxonomic practices and species characteristics, which vary across taxonomic groups and geographic regions [47]. For example, *Eschweilera coriacea* (DC.) S.A.Mori (Lecythidaceae), another hyperdominant species, has undergone several lumping events and recent molecular analysis indicates that *E. coriacea* is indeed a single species [9,70].

### Way forward: bridging taxonomy and macroecology

Our study uncovers temporal trends in the taxonomy of Amazonian palms, that is, the change in the number of accepted species across, time, regions, and genera following taxonomic revisions. For palms, the impact of revisions is substantial. However, the identified trends may not apply to other plant groups, as rates of species description and synonymization are uneven across regions and genera. To our knowledge, no studies have tracked timing of taxonomic lumping for other plant taxa, leaving it unclear how an evolving species taxonomy affects broad-scale patterns of plant richness and composition in Amazonia. This shortcoming has implications for conservation analyses. If lumping is prevalent, it may reduce species rarity [9], endemism [10], and floristic turnover [71]. Conversely, frequent species discoveries and taxonomic splitting may increase these parameters [72]. The extent to which opposing trends balance out across space and time remains unquantified for a large number of taxa [73].

Many tropical plants lack recent taxonomic revisions [69], and a small proportion of new species descriptions are supported by molecular data [74]. Thus, hidden synonyms and cryptic species will continue to bias broad-scale, multi-species ecological studies for the decades to come. This bias cannot be completely eliminated but, as we show here, it can be measured, thereby providing a baseline reference to quantify uncertainty in biogeographical, evolutionary, and macroecological analyses. The emergence of nomenclatural data infrastructures offers promising avenues for addressing this challenges.

A growing number of data infrastructure supports handling complex taxonomic data [75]. These infrastructures enables plant ecologists to explore data ontologies and taxonomic terminology to map when and how often species limits have been redefined [76]. A significant challenge is that recognizing heterotypic synonyms — unlike describing new species or designating homotypic synonyms — is not a mandatory nomenclatural act. This means that the way and extent to which heterotypic synonyms is documented in the taxonomic literature depends on the reporting practice of each individual taxonomist [58]. Consequently, it non-trivial to determine, across many taxa, when lumping occurred and how it influences species counts and identity. Undertaking this task not only requires close collaboration between ecologists and taxonomists but also calls for publishing or converting taxonomic literature in a machine-readable format. Yet, several taxonomic journals still publish articles only as portable document format (PDF), which demands substantial effort to make key information, such as synonymizations, easy to reuse [77]. Recently developed tools and services (e.g., Plazi [77] and PlutoF [78]) start to streamline this process. While their current application is still modest, they represent an important step towards a broader use of taxonomic information [79].

## Concluding remarks

Taxonomic progress is often thought of as a steady process of accumulating newly described species. However, the case of Amazonian palms suggests this view captures only part of the picture. We find that a few single revision events between the 1990s and 2000s have caused an abrupt reduction in species numbers, with uneven effects across genera and regions. Importantly, around 800 heterotypic synonyms remained undetected for up to two centuries. Our study suggests that variability in taxonomic revisions, in terms of timing and effect, may be observed for other plant groups as well. As taxonomy affects the richness, abundance, and geographical range of species, its impact should be considered in macroecological models and conservation planning. Our study highlights how much taxonomic revisions can affect our understanding of biodiversity. Without them, we risk building our ecological analyses and conservation assessments on shaky ground.

## AUTHOR’S CONTRIBUTION

**JS**: Conceptualization (lead); Data collection and curation (lead); Analysis (lead); Writing – review and editing (lead), funding acquisition (lead). **ASSP**: Conceptualization (supporting); Data collection and curation (lead); Analysis (supporting); Writing, review, and editing (supporting). **TE**: Conceptualization (supporting); Data collection and curation (supporting); Analysis (supporting); Writing – review and editing (supporting), funding acquisition (supporting). **LM**: Conceptualization (supporting); Data collection and curation (supporting); Analysis (supporting); Writing – review and editing (supporting). **RT**: Conceptualization (supporting); Analysis (supporting); Writing – review and editing (supporting): **FAM**: Conceptualization (supporting); Writing – review and editing (supporting). **RL**: Conceptualization (supporting); Data collection and curation (supporting); Analysis (supporting); Writing – review and editing (supporting). **JH**: Conceptualization (supporting); Data collection and curation (supporting); Analysis (supporting); Writing – review and editing (supporting), funding acquisition (supporting).

## ACKNOWLEDGMENTS

We thank the library staff of the Museo Nacional de Ciencias Naturales (MNCN-CSIC, Madrid) for facilitating access to taxonomic literature. We are also grateful to Cristina Ronquillo for assistance in standardizing species authorities’ names and to Sylvia Mota de Oliveira, Toby Pennington, and Martin Weiss for their comments on earlier versions of this manuscript.

## DATA ACCESSIBILITY STATEMENT

Data are publicly available at 10.6084/m9.figshare.28344428.v1

## REFERENCES

1. Kew RBG. 2016 The state of the world’s plants report. R. Bot. Gard. Kew, 1–80.

2. Edie SM, Smits PD, Jablonski D. 2017 Probabilistic models of species discovery and biodiversity comparisons. Proc. Natl. Acad. Sci. 114, 3666–3671. (doi:10.1073/pnas.1616355114)

3. er Steege H, Mota de Oliveira S, Pitman NCA, Sabatier D, Antonelli A, Guevara Andino JE, Aymard GA, Salomão RP. 2019 Towards a dynamic list of Amazonian tree species. Sci. Rep. 9, 3501. (doi:10.1038/s41598-019-40101-y)

4. Stropp J, Ladle RJ, Emilio T, Lessa T, Hortal J. 2022 Taxonomic uncertainty and the challenge of estimating global species richness. J. Biogeogr. 49, 1654–1656. (doi:10.1111/jbi.14463)

5. Knapp S, Lughadha EN, Paton A. 2005 Taxonomic inflation, species concepts and global species lists. Trends Ecol. Evol. 1, 7–8.

6. Paton AJ, Brummitt N, Govaerts R, Harman K, Hinchcliffe S, Allkin B, Lughadha EN. 2008 Towards Target 1 of the Global Strategy for Plant Conservation: A Working List of All Known Plant Species-Progress and Prospects. Taxon 57, 602–611.

7. Führding-Potschkat P, Weigelt P, Kreft H, Ickert-Bond SM. 2023 Drivers of Variation in Synonym Numbers of Angiosperm Species Names. Authorea Prepr.

8. Alroy J. 2002 How many named species are valid? Proc. Natl. Acad. Sci. 99, 3706LP–3711. (doi:10.1073/pnas.062691099)

9. Bacon CD, Hill A, Ter Steege H, Antonelli A, Damasco G. 2022 The impact of species complexes on tree abundance patterns in Amazonia. Am. J. Bot. 109, 1525–1528. (doi:10.1002/ajb2.16069)

10. Daru BH, Farooq H, Antonelli A, Faurby S. 2020 Endemism patterns are scale dependent. Nat. Commun. 11, 2115. (doi:10.1038/s41467-020-15921-6)

11. Gemeinholzer B et al. 2020 Data storage and data re-use in taxonomy—the need for improved storage and accessibility of heterogeneous data. Org. Divers. Evol. 20, 1–8. (doi:10.1007/s13127-019-00428-w)

12. POWO. 2022 Plants of the World Online. Facilitated by the Royal Botanic Gardens, Kew. See http://www.plantsoftheworldonline.org/.

13. Walker B. 2022 kewr. See https://github.com/barnabywalker/kewr.

14. Baker WJ, Dransfield J. 2016 Beyond Genera Palmarum: progress and prospects in palm systematics. Bot. J. Linn. Soc. 182, 207–233.

15. er Steege H et al. 2013 Hyperdominance in the Amazonian tree flora. Science (80-.). 342, 1243092. (doi:10.1126/science.1243092)

16. Stropp J, Umbelino B, Correia RA, Campos-Silva J V, Ladle RJ, Malhado ACM. 2020 The ghosts of forests past and future: deforestation and botanical sampling in the Brazilian Amazon. Ecography (Cop.)., 1–11. (doi:10.1111/ecog.05026)

17. RAISG. 2012 Amazônia sob Pressão., 69. See www.raisg.socioambiental.org.

18. Henderson AJ. 1995 The palms of the Amazon. Oxford University Press; New York. See http://lib.ugent.be/catalog/rug01:000386429.

19. Cardoso D et al. 2017 Amazon plant diversity revealed by a taxonomically verified species list. Proc. Natl. Acad. Sci. 114, 10695–10700.

20. Flora e Funga do Brasil 2020. 2024 Flora e Funga do Brasil 2020. Jardim Botânico do Rio de Janeiro. See http://floradobrasil.jbrj.gov.br/.

21. Tropicos. 2024 Missouri Botanical Garden. See https://tropicos.org.

22. POWO. 2024 Plants of the World Online. See https://powo.science.kew.org/.

23. WFO. 2024 World Flora Online. See http://www.worldfloraonline.org/.

24. Henderson A, Galeano G, Bernal R. 1995 Field guide to the palms of the Americas. Princeton University Press.

25. PALMWeb. In press. Palmweb: Palms of the World Online. See https://palmweb.org/ (accessed on 20 September 2022).

26. Balslev H, Kahn F, Millan B, Svenning J-C, Kristiansen T, Borchsenius F, Pedersen D, Eiserhardt WL. 2011 Species Diversity and Growth Forms in Tropical American Palm Communities. Bot. Rev. 77, 381–425. (doi:10.1007/s12229-011-9084-x)

27. IPNI. 2022 International Plant Names Index. R. Bot. Gard. Kew, Harvard Univ. Herb. Libr. Aust. Natl. Bot. Gard. See https://www.ipni.org/.

28. Turland NJ et al. 2018 International Code of Nomenclature for algae, fungi, and plants (Shenzhen Code) adopted by the Nineteenth International Botanical Congress Shenzhen, China, July 2017. Koeltz botanical books.

29. Linné C von. 1781 Supplementum plantarum Systematis vegetabilium editionis decimae tertiae, Generum plantarum editionis sextae, et Specierum plantarum editionis secunda.

30. Dugand A. 1941 Notas sobre palmas colombianas y una del Brasil. Caldasia 1, 17–29.

31. Wessels Boer JG. 1968 The geonomoid palms. Meded. van het Bot. Museum en Herb. van Rijksuniv. te Utr. 282, 1–202.

32. Henderson AJ. 2020 A revision of Attalea (Arecaceae, Arecoideae, Cocoseae, Attaleinae). Phytotaxa 444, 1–76. (doi:10.11646/phytotaxa.444.1.1)

33. Burret (Maximilian) Karl Ewald. 1929 Die Palmengattungen Orbignya, Attalea, Scheelea und Maximiliana. Notizblatt des Bot. Gartens und Museums zu Berlin-Dahlem 10.

34. Bartlett HH. 1935 Scheelea Lundellii, a New” corozo” Palm from the Department of Petén, Guatemala. Carnegie Inst. Wash.

35. Bailey LH. 1947 Indigenous palms of Trinidad and Tobago. Gentes Herb.

36. GBIF. 2022 GBIF Occurrence Download 10.15468/dl.m8jkhk. See 10.15468/dl.m8jkhk.

37. Chamberlain S, Ram K, Barve V, Mcglinn D, Chamberlain MS. 2017 Package ‘rgbif’. Interface to Glob. Biodivers. Inf. Facil. ‘API 5.

38. Ribeiro BR, Velazco SJE, Guidoni-Martins K, Tessarolo G, Jardim L, Bachman SP, Loyola R. 2022 bdc: A toolkit for standardizing, integrating and cleaning biodiversity data. Methods Ecol. Evol. 13, 1421–1428.

39. Hosmer DW, Lemeshow S, May S. 2008 Applied survival analysis: regression modeling of time-to-event data. John Wiley & Sons.

40. Therneau TM. 2024 A package for survival analysis in R.

41. Alboukadel K, Kosinski M, Przemyslaw B, Scheipl F. 2022 Package ‘survminer’.

42. Bates D. 2010 lme4: Linear mixed-effects models using S4 classes. http://cran.r-project.org/web/packages/lme4/index.html

43. Onstein RE, Baker WJ, Couvreur TLP, Faurby S, Svenning J-C, Kissling WD. 2017 Frugivory-related traits promote speciation of tropical palms. Nat. Ecol. Evol. 1, 1903–1911.

44. Maitner BS et al. 2018 The bien r package: A tool to access the Botanical Information and Ecology Network (BIEN) database. Methods Ecol. Evol. 9, 373–379.

45. Rivas-Chamorro M, Cadenillas R, Ge X-J, Jin L, Millán B, Roncal J. 2023 Testing species relationships and delimitation in the Amazonian hyperdominant Astrocaryum section Huicungo (Arecaceae) using chloroplast data from genome skimming. Taxon 72, 501–514. (doi:10.1002/tax.12928)

46. Paola de Lima Ferreira; Angela Cano; Benedikt Kuhnhuser; Sidonie Bellot; William Baker; Wolf Eiserhardt; the Palm Phylogeny Working Group. 2024 Palms in space and time: Progress towards a genomic species-level phylogeny. In XX International Botanical Congress, Madrid.

47. Padial JM, De la Riva I. 2021 A paradigm shift in our view of species drives current trends in biological classification. Biol. Rev. 96, 731–751.

48. Mori SA. 1992 Neotropical Floristics and Inventory: Who Will do the Work? Brittonia 44, 372–375. (doi:10.2307/2806943)

49. Henderson AJ. 2011 A revision of Desmoncus (Arecaceae). Phytotaxa 35, 1–88.

50. Henderson AJ. 2000 Bactris (Palmae). Flora Neotrop. 79, 1–181.

51. Henderson AJ. 2011 A revision of Geonoma (Arecaceae). Revisión de Geonoma (Arecaceae). Phytotaxa, 1–271.

52. Henderson A. 2011 A revision of Geonoma (Arecaceae). Phytotaxa 17, 1–271.

53. Maas PJM, Westra LYT, Arias Guerrero S, Lobão A, Scharf U, Zamora NA, Erkens R. 2015 Confronting a morphological nightmare: Revision of the Neotropical genus Guatteria (Annonaceae). Blumea J. plant Taxon. plant Geogr. 60, 1–219. (doi:10.3767/000651915X690341)

54. Lucas EJ, Matsumoto K, Harris SA, Nic Lughadha EM, Benardini B, Chase MW. 2011 Phylogenetics, morphology, and evolution of the large genus Myrcia sl (Myrtaceae). Int. J. Plant Sci. 172, 915–934.

55. Lima DF, Goldenberg R, Lucas E. 2018 Taxonomic novelties in Myrcia guianensis and allied species (Myrtaceae: Myrteae), including mass-typification in a large and taxonomically challenging group. Kew Bull. 73, 1–17.

56. Christenhusz MJM, Byng JW. 2016 The number of known plants species in the world and its annual increase. Phytotaxa 261, 201–217.

57. Antonelli A et al. 2020 State of the World’s Plants and Fungi.

58. Lughadha EN, Govaerts R, Belyaeva I, Black N, Lindon H, Allkin R, Magill RE, Nicolson N. 2016 Counting counts: revised estimates of numbers of accepted species of flowering plants, seed plants, vascular plants and land plants with a review of other recent estimates. Phytotaxa 272, 82–88.

59. Wortley AH, Scotland RW. 2004 Synonymy, Sampling and Seed Plant Numbers. Taxon 53, 478–480. (doi:10.2307/4135625)

60. Moonlight PW, Baldaszti L, Cardoso D, Elliott A, Särkinen T, Knapp S. 2024 Twenty years of big plant genera. Proc. R. Soc. B Biol. Sci. 291, 20240702. (doi:10.1098/rspb.2024.0702)

61. West-Eberhard MJ. 2003 Developmental plasticity and evolution. Oxford University Press.

62. Weber UK, Nuismer SL, Espíndola A. 2020 Patterns of floral morphology in relation to climate and floral visitors. Ann. Bot. 125, 433–445. (doi:10.1093/aob/mcz172)

63. Figueiredo FOG et al. 2022 Linking high diversification rates of rapidly growing Amazonian plants to geophysical landscape transformations promoted by Andean uplift. Bot. J. Linn. Soc. 199, 36–52. (doi:10.1093/botlinnean/boab097)

64. Svenning J-C, Borchsenius F, Bjorholm S, Balslev H. 2008 High tropical net diversification drives the New World latitudinal gradient in palm (Arecaceae) species richness. J. Biogeogr. 35, 394–406. (doi:10.1111/j.1365-2699.2007.01841.x)

65. Kissling WD, Baker WJ, Balslev H, Barfod AS, Borchsenius F, Dransfield J, Govaerts R, Svenning J-C. 2012 Quaternary and pre-Quaternary historical legacies in the global distribution of a major tropical plant lineage. Glob. Ecol. Biogeogr. 21, 909–921. (doi:10.1111/j.1466-8238.2011.00728.x)

66. Gomes VHF, Vieira ICG, Salomão RP, ter Steege H. 2019 Amazonian tree species threatened by deforestation and climate change. Nat. Clim. Chang. 9, 547–553. (doi:10.1038/s41558-019-0500-2)

67. Alvez-Valles CM, Balslev H, Garcia-Villacorta R, Carvalho FA, Menini L. 2018 Palm species richness, latitudinal gradients, sampling effort, and deforestation in the Amazon region. Acta Bot. Brasilica 32, 527–539.

68. Hopkins MJG. 2007 Modelling the known and unknown plant biodiversity of the Amazon Basin. J. Biogeogr. 34, 1400–1411.

69. Grace OM et al. 2021 Botanical Monography in the Anthropocene. Trends Plant Sci. 26, 433–441. (doi:10.1016/j.tplants.2020.12.018)

70. Heuertz M et al. 2020 The hyperdominant tropical tree Eschweilera coriacea (Lecythidaceae) shows higher genetic heterogeneity than sympatric Eschweilera species in French Guiana. Plant Ecol. Evol. 153, 67–81.

71. Tuomisto H. 2010 A diversity of beta diversities: straightening up a concept gone awry. Part 1. Defining beta diversity as a function of alpha and gamma diversity. Ecography (Cop.). 33, 2–22. (doi:10.1111/j.1600-0587.2009.05880.x)

72. Damasco G, Baraloto C, Vicentini A, Daly DC, Baldwin BG, Fine PVA. 2021 Revisiting the hyperdominance of Neotropical tree species under a taxonomic, functional and evolutionary perspective. Sci. Rep. 11, 9585. (doi:10.1038/s41598-021-88417-y)

73. Lessa T, Stropp J, Hortal J, Ladle RJ. 2024 How taxonomic change influences forecasts of the Linnean shortfall (and what we can do about it)? J. Biogeogr. 51, 1365–1373. (doi:10.1111/jbi.14829)

74. Miralles A et al. 2020 Repositories for Taxonomic Data: Where We Are and What is Missing. Syst. Biol. 69, 1231–1253. (doi:10.1093/sysbio/syaa026)

75. Tuominen J, Laurenne N, Hyvönen E. 2011 Biological names and taxonomies on the semantic web–managing the change in scientific conception. In Extended Semantic Web Conference, pp. 255–269. Springer.

76. Franz NM, Peet RK. 2009 Towards a language for mapping relationships among taxonomic concepts. Syst. Biodivers. 7, 5–20.

77. Agosti D, Catapano T, Sautter G, Egloff W. 2019 The Plazi Workflow: The PDF prison break for biodiversity data. Biodivers. Inf. Sci. Stand.

78. Abarenkov K et al. 2010 PlutoF—a web based workbench for ecological and taxonomic research, with an online implementation for fungal ITS sequences. Evol. Bioinforma. 6, EBO–S6271.

79. Upham NS et al. 2021 Liberating host–virus knowledge from biological dark data. Lancet Planet. Heal. 5, e746–e750.

